# Learning the sequence code of protein expression in human immune cells

**DOI:** 10.1101/2023.09.01.555843

**Authors:** Benoît P. Nicolet, Anouk P. Jurgens, Kaspar Bresser, Aurélie Guislain, Antonia Bradariç, Monika C. Wolkers

## Abstract

Accurate protein expression in human immune cells is essential for appropriate cellular function. The mechanisms that define protein abundance are complex and executed on transcriptional, post-transcriptional and post-translational level. Here, we present SONAR, a machine learning pipeline that learns the endogenous sequence code and that defines protein abundance in human cells. SONAR uses thousands of sequence features (SFs) to predict up to 63% of the protein abundance independently of promoter or enhancer information. SONAR uncovered the cell type-specific and activation-dependent usage of SFs. The deep knowledge of SONAR provides a map of biologically active SFs, which can be leveraged to manipulate the amplitude, timing, and cell type-specificity of protein expression. SONAR informed on the design of enhancer sequences to boost T cell receptor expression and to potentiate T cell function. Beyond providing fundamental insights in the regulation of protein expression, our study thus offers novel means to improve therapeutic and biotechnology applications.

**One Sentence Summary:** SONAR informs the design of cell type-specific protein expression in human cells

## INTRODUCTION

The development, function and long-term maintenance of immune cells requires an intricate regulation of protein expression, which is defined by the interplay of transcriptional, post-transcriptional and post-translational events (*1*). Understanding the rules that control the protein expression in any given cell type, and that capture alterations of protein abundance upon a cell receiving external signal is - beyond fundamental interest - critical to improve and develop novel therapeutic interventions.

mRNA expression levels depend on a crosstalk of sequence features (SFs). This includes promoter motifs in the DNA with their respective interactors, such as transcription factors, which constitute a key factor for defining RNA expression levels (*2*, *3*). In addition, many hard-wired features of the mRNA define the protein output and thus the protein abundance (*4*, *5*). These features can be broadly separated into 2 groups: static regulators, such as the GC-content and sequence conservation (*4*, *6*–*8*), and dynamic regulators, such as mRNA interaction with RNA-binding proteins (RBPs), some of which lead to mRNA modifications (e.g. m6A), or interaction with microRNAs (miRs; (*9*, *10*)). Other dynamic regulators for protein abundance include the tRNA abundance (*11*), and post-translational modifications (PTMs) of amino acids residues (e.g. ubiquitination of lysines (*12*)). Combined, these regulatory features imprint the cell’s identity, and they allow it to respond to environmental cues. Immune cells are particularly well adapted to swiftly alter their phenotype, which prepares them for immunological insults. They do so by controlling for example mRNA decay and translation (*13*–*15*). Despite our current knowledge that all of the aforementioned regulatory processes contribute to protein abundance, a comprehensive map of their relative contribution to this key feature of cellular identity and function is lacking.

Machine learning (ML) has been instrumental to reveal many aspects of gene expression (*16*), such as the relation of sequence characteristics in transcriptional regulation of mammalian cells (*17*) or its gene regulatory structure (*18*), codon-usage association with mRNA stability in yeast (*19*) and the decay rates in mammalian cells (*20*). While instrumental to our understanding, these foundational studies focus on one aspect of the complex protein expression regulation and are yet to be integrated to reveal the relative contribution of sequence feature to protein abundance.

In this study, we developed SONAR, an ML pipeline that allowed us to 1) learn the endogenous, cell-type specific sequence code that defines the protein abundance in immune cells; 2) map the relative contribution of SFs to protein abundance under steady state conditions, and upon T cell activation; and 3) manipulate the amplitude, and timing of protein expression in human cells. Lastly, we leveraged the knowledge captured by SONAR to enhance protein expression in T cells, resulting in enhanced immune capacity.

## RESULTS

### SONAR captures the sequence feature code for protein expression in human cells

To predict protein abundance, SONAR relies on a SF library on the entire human proteome. To generate the SF library, we aggregated mRNA-level SFs for the entire protein-coding human transcriptome (94,348 mRNA isoforms, see Methods). SFs included “static” SFs such as GC content and sequence conservation, and “dynamic” SFs such as codon usage (influenced by the dynamic tRNA abundance), predicted miR seed score (miRDB), RBP motifs (ATracT), and predicted mRNA modification sites (RMVar). We combined mRNA-level features identified by their respective location (5’UTR, CDS, 3’UTR, and full-length mRNA) with protein level information such as amino-acid usage and predicted protein modification sites (PTMdb) resulting in a total of 7,225 SFs for each protein of the human proteome (Fig. 1A, S1A).

**Fig. 1:**
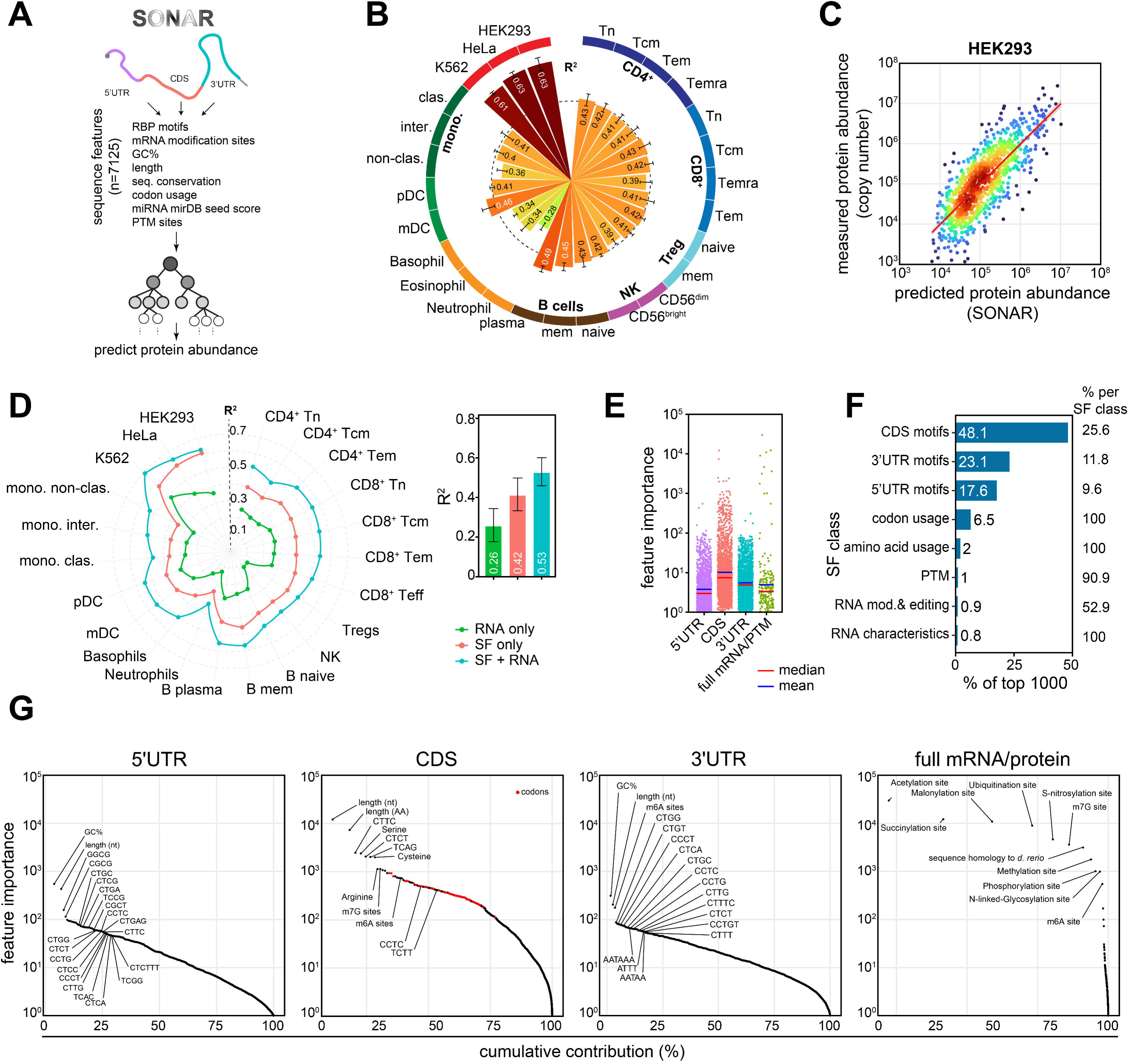
Modelling the sequence code of protein abundance. (**A**) SONAR was trained to learn the code for protein abundance using sequence features (SFs) found in the human transcriptome (*see Methods*, Fig S1A,B). (**B-C**) Protein abundance model performance on test sets (R^2^, indicated in bar, with S.D.) in (B) indicated human cell types (n=121; 2-4 biological replicates; dashed line shows mean R^2^), or (C) HEK293 cells (R^2^: 0.618; representative of 3; red line shows linear regression line). (**D**) Protein abundance model performance for all cell types (left panel) trained on mRNA abundance data (RNA only), sequence features (SF only) or both (SF+RNA). Average performance per model type (right panel; n=22; average is indicated in bar; with S.D.). (**E**) SF contribution per mRNA region of origin (feature importance (a.u.)). (**F**) Prevalence of SF class within the top 1000 most predictive SFs. Percentages (of total) per SF type are indicated on the right. (**G**) Cumulative contribution of all SFs split per region of origin. Codons are indicated in red. Tn: naïve, Tcm: central-memory, Tem: effector memory, Teff: effector T cells; Treg: regulatory T cells; mem: memory; pDC: plasmacytoid, mDC: myeloid dendritic cells; mono.: monocytes; clas.: classical; inter.: intermediate; NK: natural killer cells.

We next trained SONAR to predict protein abundance based on SFs alone (S1B). We used published protein abundance data of three different cell lines (n=9; (*21*); Fig. 1B) and a wide variety of peripheral blood-derived human immune cell subsets (n=112; (*22*)); Table S1). To prevent data leakage, we held out 20% of the data to test the models, and we used the remaining 80% of the data for training. Models trained on randomly shuffled protein abundance data lacked any predictive capacity (R^2^: 0.0008). Solely relying on SFs, SONAR accurately predicted 40.7% of the protein abundance in primary immune cells and up to 62.7% in cell lines (Fig. 1B, C). This finding indicates that SFs contain information that are important for predicting protein abundance.

Notably, SONAR was superior (R^2^:0.42) to experimental mRNA measurements alone (R^2^:0.26) in predicting protein abundance (Fig. 1D). However, when we included mRNA expression data to SONAR models (SF+RNA), the overall prediction accuracy improved even further by on average 11% (R^2^:0.53, Fig. 1D, blue data points). SF+RNA models reached up to 69% in cell lines (Fig. 1D). This finding indicates that mRNA abundance rules are already partially captured by SONAR, yet mRNA abundance remains a critical data modality to improve the prediction of protein abundance from SFs alone.

SONAR uses XGBoost (*23*), which generates auditable models. It can reveal the relative contribution of individual SFs by quantifying the effect of the loss of a given feature to the protein abundance models. This analysis revealed that not all SFs contributed equally to SONAR’s predictive capacity. In fact, 2243 SFs (33.06% of all SFs) are required to reach 95% of the model’s full predictive capacity when using SFs alone (Fig. S1C; Data S1). Notably, even though the 5’UTR and 3’UTR are considered the prime regions regulating the protein abundance (e.g. by translation control), SONAR uncovered that the importance of SFs from the CDS is on average higher (Fig. 1E, F). This finding corroborates with the strong regulatory capacity reported for the CDS of the tumor repressor gene TP53/p53 (*24*), and the high level binding of RBPs to the CDS in cell lines (*25*). Combined, these findings argue that SFs within the CDS exert a high potential regulatory role, which is at least on par with that of the UTRs.

We next analyzed the individual SF contribution to protein abundance. The most predictive SFs were dominated by sequence motifs, such as GC and TOP-like CT-rich motifs (*26*) located in the 5’UTR and the CDS, as well as CT-rich, AU-rich motifs and canonical poly-adenylation signal sites in the 3’UTR (Fig. 1G). Other top-contributing SFs were GC content, region length, codon usage in the CDS, and putative m6A and m7G sites in the CDS, the 3’UTR, and the full length mRNA models. Also putative PTM sites such as acetylation and ubiquitination sites displayed high SF importance (Fig. 1G). Intriguingly, SONAR also revealed position-dependent SF effects, as exemplified by the effect of m6A predicted sites within different mRNA regions (Fig. S2). Combined, SONAR can model, understand, and predict the SF landscape associated with protein abundance.

### The contribution of sequence features is cell type-specific

Provided that mRNA regulators such as miRNAs and RBPs display a cell-type specific expression (*27*, *28*), we next assessed how SF usage varies across cell types. UMAP clustering and comparison of SF importance in the SONAR models between cell types revealed a distribution that closely resembled the biological origin of cell types (Fig. 2A, Fig. S3, Data S2). Furthermore, cells with very similar gene expression and cell state, i.e. CD4^+^ and CD8^+^ naïve T cells, are more akin in their SF usage than when compared to i.e. HeLa cells (Fig. 2B), suggesting that the regulatory landscape of HeLa cells differs from that of T cells. To further characterize the variability of SF usage, we isolated SFs with low (termed ‘core’) or high variability in SF importance across all cell types. SFs in the CDS were not only contributing most to protein abundance models (Fig 1E). They were also the most stable across cell types (Fig. 2C). Conversely, SFs in the UTRs were also important but displayed a much higher inter-cell variability (Fig. S3). Combined, this analysis indicates that the SF landscape is cell-type specific.

**Fig. 2:**
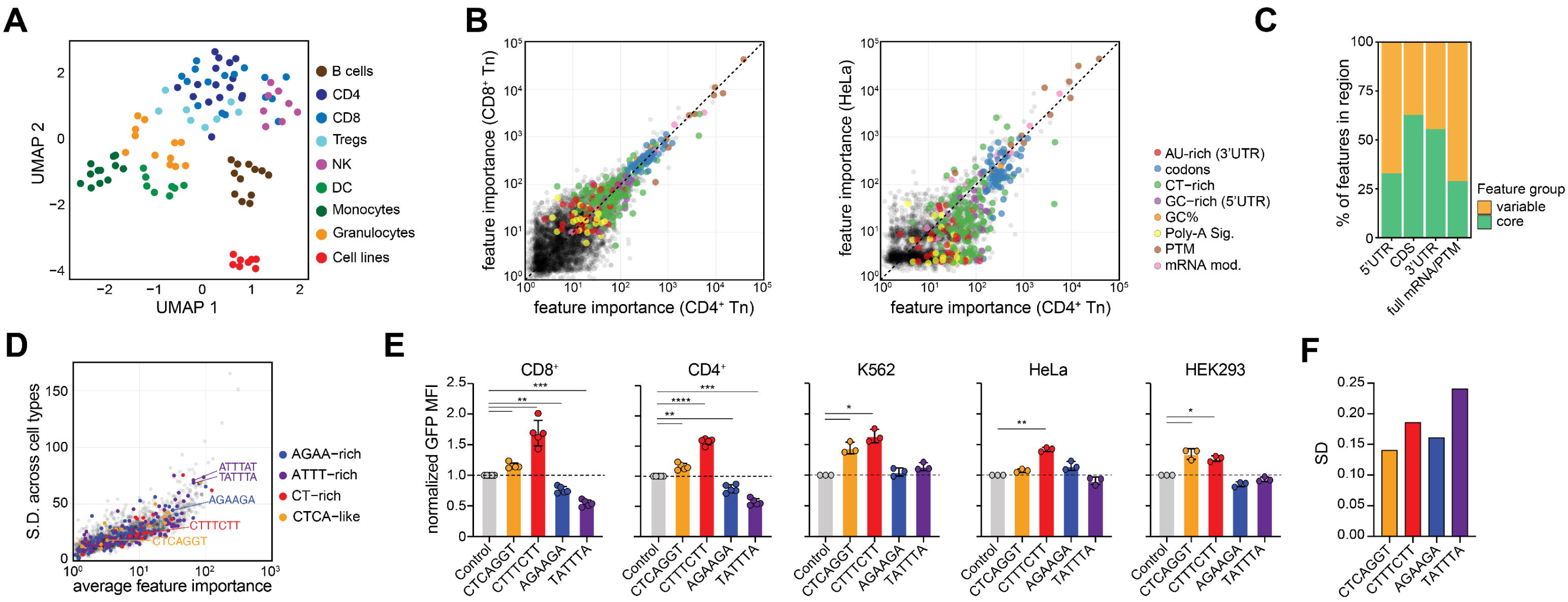
The sequence code of protein abundance is cell type specific. (**A**) UMAP representation of feature importance usage per sample in protein abundance models of Fig 1. (**B**) Comparison of the average SF importance in protein abundance models of CD4^+^ with CD8^+^ naïve T cells or with HeLa cells, with indicated color-coded SF types. (**C**) Variability of SF importance across cell types grouped into ‘core’ (low variability) or ‘variable’ across cell (see methods; supplemental note #1) represented as percentage per feature groups. (**D**) Comparison of the average SF importance and respective S.D. across all cell types for 3’UTR SFs. Color depicts the indicated motif family. (**E**) Syn3UTR-GFP reporter assay for the indicated 3’UTR motifs (see methods). GFP expression levels determined by flow-cytometry and normalized to scrambled (control) 3’UTR in indicated cell types (n=3-5; ANOVA with post-hoc Tukey test). (**F**) S.D. across cell types for indicated 3’UTR motifs.

To experimentally validate the cell-type specificity of SFs, we selected four 3’UTR motifs with varying variability (S.D. across cell types) and SF importance (Fig. 2D). We generated a 180nt-long synthetic 3’UTR sequence (syn3UTR) devoid of important SF features in which we inserted 6 occurrences of each motif individually (Data S3). The obtained syn3UTRs were fused to the 3’end of a GFP reporter gene and introduced into 5 different cell types (Fig. 2E, Fig. S4). All motifs had some level of cell type specificity, but to a different degree. For example, CTTTCTT-containing 3’UTR reporters consistently boosted the GFP protein expression compared to scrambled syn3UTR control in T cells and all tested cell lines (Fig. 2E). In contrast, TATTTA repressed GFP expression most effectively in T cells (Fig. 2E). Intriguingly, the variability (S.D.) of the experimental data points per motif across all cell types resembled the variability of SF importance that was predicted by SONAR (Fig. 2D-F). Combined, SONAR models identify both core and cell-type specific SF usage for regulating protein abundance.

### CT-rich 3’UTR motifs enhance protein expression

The design of synthetic 3’UTRs can manipulate protein expression without altering the coding sequence. To experimentally validate whether a high feature importance that was assigned by SONAR correctly predicts biological significance, we devised a massively parallel reporter assay (MPRA) of syn3UTR containing 6 occurrences of 6-mer or 7-mer motifs (n=467 motifs; Data S4). The resulting syn3UTR library was introduced into HEK293 cells and into CD4^+^ and CD8^+^ T cells, and the syn3UTR enrichment was calculated between GFP^hi^ cells and GFP^lo^ FACS-sorted CD4^+^ T cells, CD8^+^ T cells, and Hek293 cells by sequencing (Fig. 3A, Fig. S5A-C; Data S4). The MPRA analysis uncovered distinct biological effects of syn3UTR motifs on GFP expression (Fig. 3B, Fig. S5D, E). Notably, SFs predicted to have a high feature importance with SONAR also displayed strong biological effects with MPRA. In particular the CT- and AG-content of SFs predicted by SONAR showed high regulatory capacity on the protein expression (Fig. 3B, C; Fig S5D, E). Furthermore, whereas SONAR correctly predicted the feature importance of SFs, the MPRA uncovered the directionality of their biological effect on protein expression: CT-rich motifs in the syn3UTR enhanced the protein expression, and AG-rich motifs were mostly repressive (Fig. 3B, C; Fig. S5D, E). These divergent effects of CT- and AG-rich motifs were observed across all three tested cell types, indicating conserved regulatory mechanisms (Fig 3D). Yet, the MPRA also uncovered the cell type specificity of SF usage (Fig. 3E). These findings confirmed the potency of SONAR in predicting protein abundance, and it uncovered the role of CT- and AG-rich 3’UTR motifs in modifying protein expression in different cell types.

**Fig. 3:**
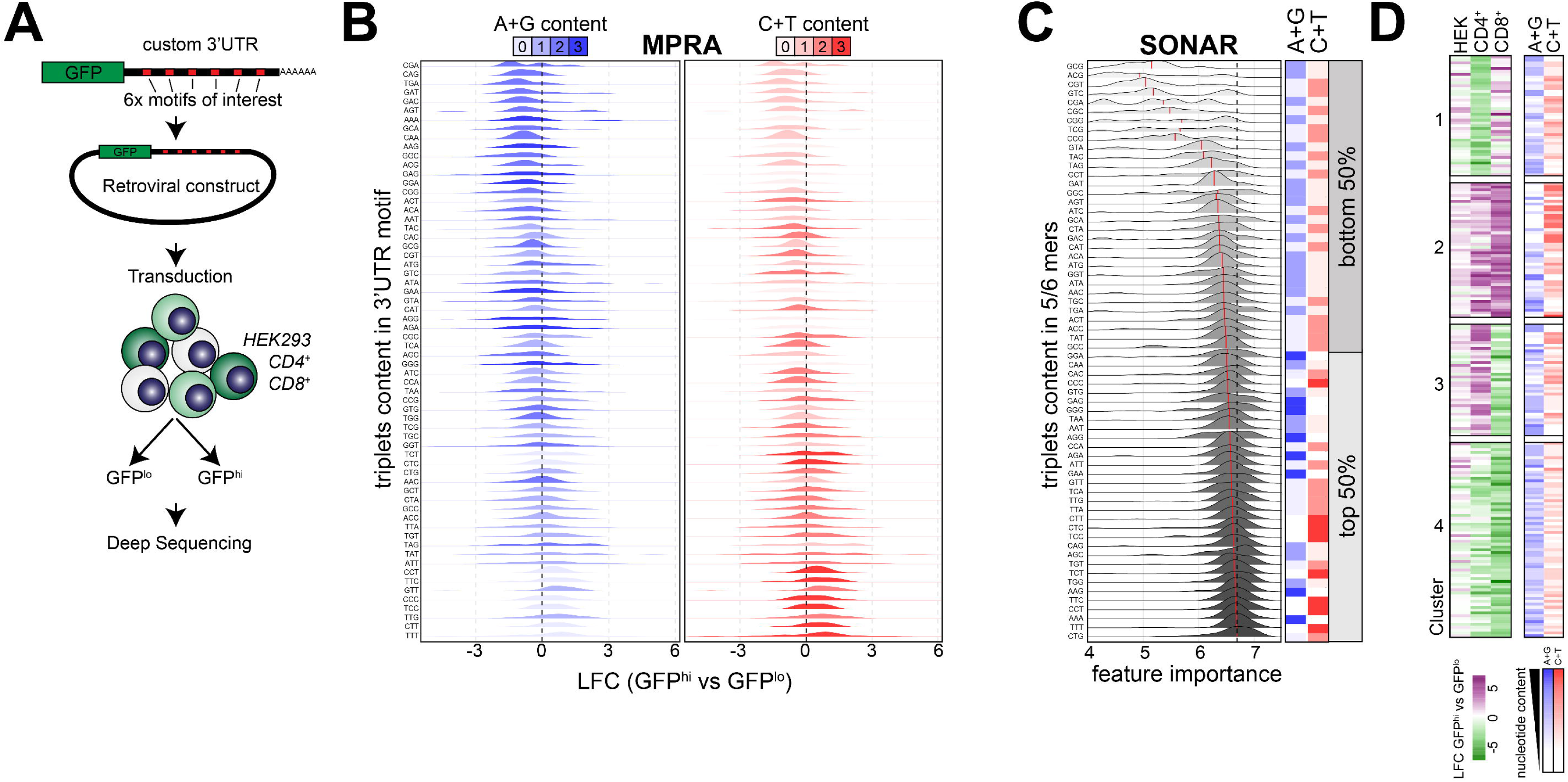
CT-rich 3’UTR motifs generally enhance protein expression. (**A**) Schematic representation of the massively parallel reporter assay (MPRA) with syn3UTR-GFP reporter assessed in CD4^+^, CD8^+^ T cells and HEK293 cells. (**B**) Log2 fold-change (LFC) enrichment of motifs found in GFP high (GFP^hi^) or low (GFP^low^) in HEK293 cells. (**C**) SONAR feature importance relation to triplet content of 5/6-mers sequence motifs found in 3’UTR derived from protein abundance models (Fig. 1). Note that feature importance does not embody the directionality of the effect. Color coding in (B, C) indicate the C+T or A+G nucleotide content of the triplet within 3’UTR motifs. The triplet content is arranged by median LFC (B) or feature importance (C). (**D**) K-means clustering (k=4) of motifs based on LFC (GFP^hi^ – GFP^lo^) in the three indicated cell types and the relation with A+G or C+T content within motifs.

### SONAR captures alterations in SF landscape upon T cell activation

We next examined whether and how SF importance for protein abundance alters when the cell state changes. T cells undergo swift and profound transcriptome and proteome remodelling upon activation and are highly dependent on post-transcriptional events for gaining T cell effector function (*13*, *14*, *29*). Therefore, T cells are an ideal model system to study the dynamics of post-transcriptional control (*7*, *23*). To uncover SF importance throughout T cell activation, we used SONAR to model the dynamic mRNA abundance, protein abundance, and the translation rate at different time points during the first 24-48h of T cell activation ((*14*); Fig. S6A; Data S5). Of note, for mRNA abundance prediction, SONAR was modified to only contain mRNA-level information (*see methods;* Fig S1A).

Interestingly, upon T cell activation, SF features most prominently contributed to all three models when located in the CDS or in the 3’UTR (Fig 4A). Furthermore, the importance of SFs substantially altered throughout activation for RNA abundance, translation, and protein abundance models (Fig 4B). Nevertheless, all models followed the same pattern in that the CDS and 3’UTR were the least variable, with about half belonging to core SFs (Fig 4B). The SF variability was the highest within the 5’UTR and the full mRNA/protein models (Fig 4B). In line with these findings, we observed considerable changes in the top-contributing SFs in all models throughout T cell activation (Fig 4C). For instance, codon usage, which can influence mRNA levels through translation-dependent decay (*19*), in addition to protein translation rates and abundance (*30*) was amongst the top contributing SFs in T cells (Fig 4C, *CDS panel*). Indeed, T cell activation was shown to induce changes in abundance of transfer RNAs (tRNA) (*31*), a feature that was well captured by SONAR. This is exemplified by the AAG codon (encoding lysine), which is known to be critical for T cell activation, and which is buffered in abundance prior to T cell activation ((*32*); Fig S6B). SONAR not only captured known relations of codon usage importance during T cell activation. It also extended this list by providing the relative contribution of all codon and amino acid SFs to mRNA, translation rate and protein abundance (Fig S6B, C). Intriguingly, codon usage aggregated per corresponding amino acid did not always show equal association with mRNA, translation rate and protein abundance. For example, the codon usage for Lysine (K) & Aspartic acid (D) had a weaker linear relationship with mRNA, yet they were positively associated with protein rate and abundance (Fig S6B, C). Thus, SONAR captures the dynamic remodelling or the SF landscape upon T cell activation.

**Fig. 4:**
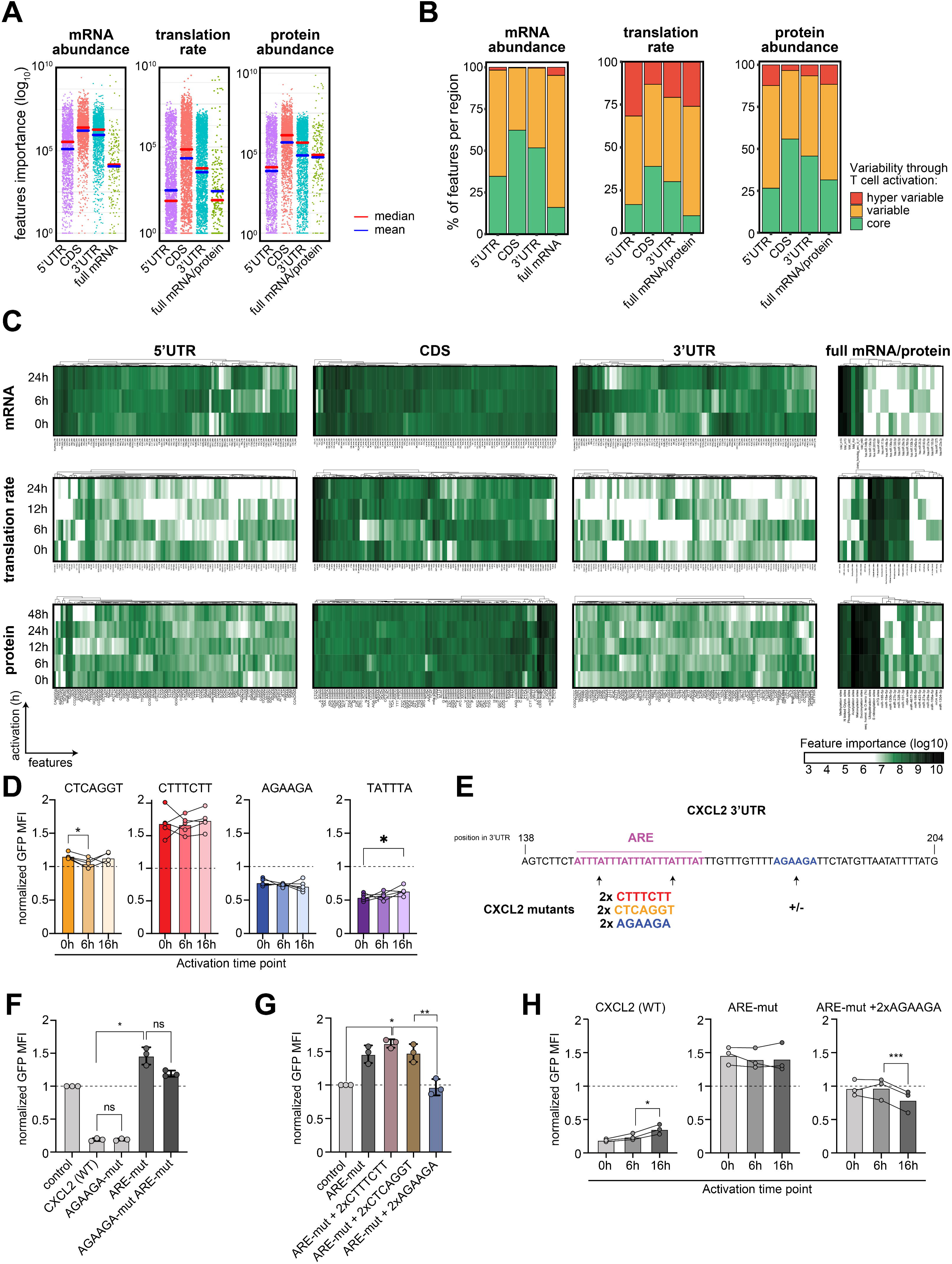
SF landscape alters upon T cell activation. (**A**) Feature importance per region of origin of mRNA abundance, translation rate, and protein abundance SONAR models of CD4^+^ T cell during activation ((*14*);average of 3 donors across timepoint). (**B**) SF importance of models in (A) was grouped by variability across time points into least variable (core), variable, and hypervariable (see methods). (**C**) Heatmap of the 100 top most predictive features per region of origin (top 25 for full mRNA/protein SFs). (**D**) Syn3UTR-GFP construct were transduced into T cell rests, and subsequently activated for 6h or 16 h with PMA-ionomycin. GFP levels were assessed by flow-cytometry and normalized to control (scrambled) at the respective timepoints. (**E**) CXCL2 3’UTR fragment (138-204nt) was mutated to replace AU-rich element (ARE) by the indicated motifs. (**F-H**) GFP levels measured by flow-cytometry of the indicated constructs at rest (F,G) or at T cell activation as in (D). (D, F-H) difference tested with ANOVA with post-hoc Tukey test. WT: wild type; mut: mutated depicts the replacement of selected motifs by scrambled padding or indicated motif.

### Sequence context defines contribution of SFs in the 3’UTR to control protein expression

To validate the alterations in SF importance observed upon T cell activation, we turned to GFP-syn3UTR constructs employed in Fig 2E, and we measured the protein expression in resting and in PMA/Ionomycin-activated T cells (Fig 4D). Protein expression under control of CTTTCTT and AGAAGA syn3UTR was insensitive to T cell activation (Fig 4D). Conversely, CTCAGGT syn3UTR induced a slight increase at 6 h of T cell activation (Fig 4D). TATTTA-containing syn3UTR expression resulted in a small but steady increase in GFP expression throughout T cell activation (Fig 4D), which is in line with previously reported relief of translation blockade by AREs in the *IFNG-*3’UTR (*13*).

We next tested the effect of 3’UTR motifs in the context of a naturally occurring 3’UTR. A CXCL2 3’UTR fragment previously used to study its regulatory capacity (*33*) contains a stretch of ARE motifs, together with one AGAAGA element (Fig. 4E, Data S3). Mutating the AGAAGA motif from the CXCL2 3’UTR did not significantly alter the regulatory capacity CXCL2 sequence (Fig 4F). However, mutating the natural ARE element fully relieved the repressive effect of the CXCL2 3’UTR to a similar extent as double depletion of AREs and the AGAAGA motif (Fig 4F). We then exchanged the ARE motif by either two CTCAGGT or CTTTCTT motifs. Intriguingly, despite the previously observed positive effect of CTTTCTT motifs on protein expression (Fig 2), this exchange did not result in further increases of GFP expression (Fig 4G). In contrast, exchanging the ARE motif with two AGAAGA motifs partially restored the suppressive activity of the CXCL3 3’UTR (Fig 4G). These finding reveals that a potentially negative sequence context can attenuate the positive effect of a 3’UTR motif (i.e. CTTTCTT), and that the sequence context contributes to the activity of SFs.

We next tested how CXCL2 3’UTR mutants modulated GFP expression upon T cell activation. The ARE-containing CXCL2 WT 3’UTR showed increased GFP expression at 16 h of T cell activation, which was abrogated when AREs were mutated (Fig 4H, (*13*)). Substitution of AREs with AGAAGA even further reduced the protein expression upon T cell activation (Fig 4H). Altogether, our findings highlight that the combination of sequence context and 3’UTR motifs defines the control of protein production. In addition, we demonstrate that 3’UTR motifs can be used to manipulate the timing of protein expression upon T cell activation.

### SONAR informs on the potential mechanism of action of SFs

Finally, we sought to define the relative contribution of SFs to mRNA abundance, translation rate, and protein abundance models throughout T cell activation. Translation rate and protein models preferentially used CDS parameters such as length, GC content and TOP-like (CT-rich) motifs (Fig. 5A). In line with previous reports (*6*), m1A, m5C, m6A, m7G putative sites strongly contributed to all 3 models, but most dominantly to mRNA abundance models (Fig. 5A). m7G modifications have been associated with translation efficiency (*34*). Intriguingly, SONAR suggests a novel role for m7G sites in regulating mRNA abundance, which was confirmed by re-analysis of transcriptome-wide internal m7G mapping data ((*34*); Fig S8; Data S6). Thus, SONAR not only maps the dynamic contribution of SFs to predict protein abundance, but it also uncovers their mechanism of action.

**Fig. 5:**
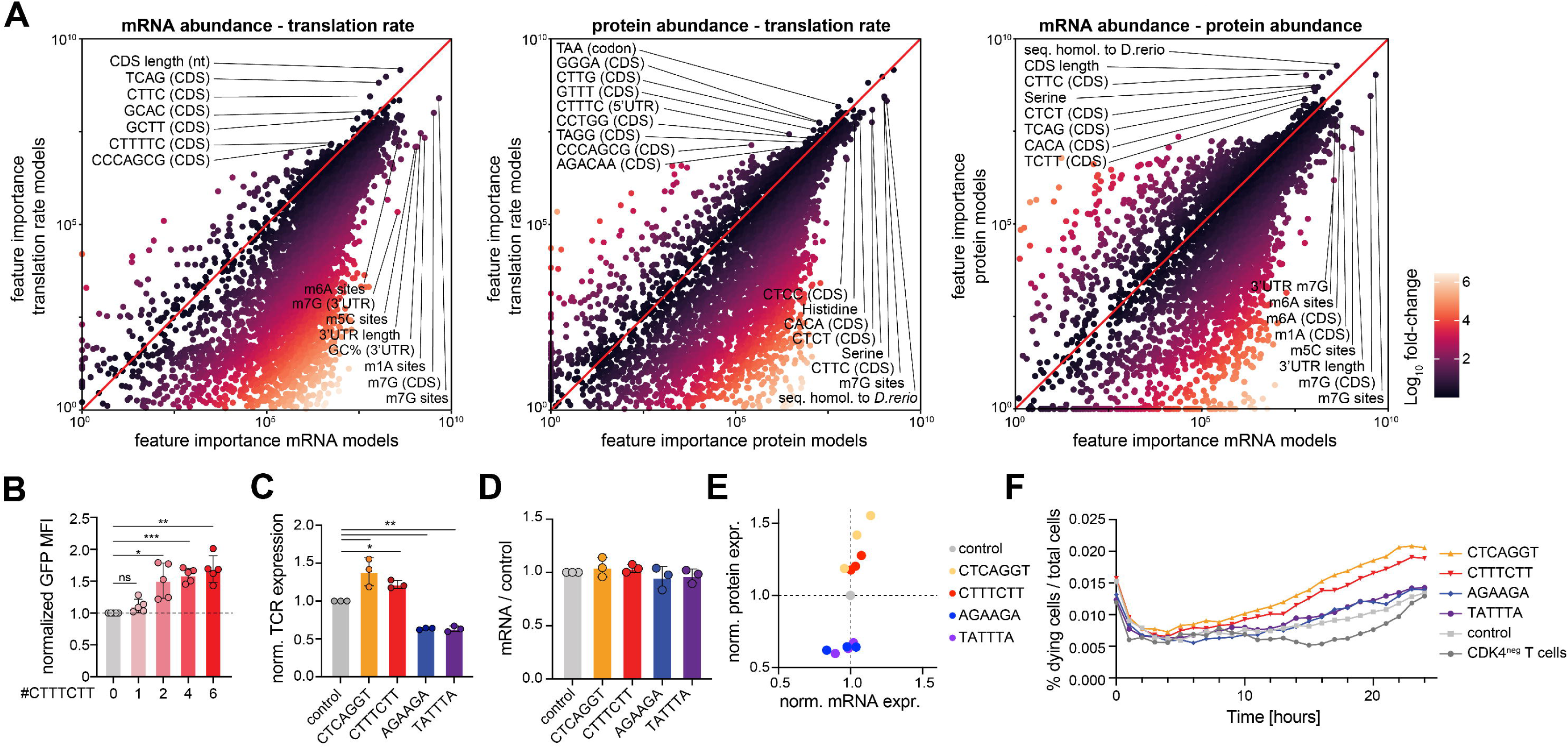
SONAR-informed synthetic 3’UTRs modulates protein expression to enhance T cell immune function. (**A**) Comparison of SF contribution in models of Fig 4A on mRNA abundance, translation rate, and protein abundance. Color represents log10-fold change. (**B**) GFP levels of syn3UTR-GFP with indicated motifs and occurrence transduced into CD8^+^ T cells. (**C,D**) syn3UTR with indicated motifs were introduced in the CDK4 transgenic T cell receptor (TCR) and TCR protein (C; flow cytometry; MFI) or mRNA expression (D; qPCR) was quantified and normalized to the control (scrambled). (**E**) Comparison of normalized mRNA and protein expression from (C,D). (**F**) FACS-sorted CD8^+^ T cells expressing the CDK4_R>L_ TCR were co-cultured for 24hours with tumor cells expressing the CDK4_R>L_ neoantigen at a 1:1. Cell death across time was quantified by caspase-3/7 apoptosis assay in IncuCyte (n=3 donors; see method). (B-D) differences assessed by ANOVA with post-hoc Tukey test.

### Boosting T cell function by 3’UTR-mediated control of translation of the T cell receptor

T cells have been successfully used for immunotherapy, for example by making use of T cell receptors (TCR) to specifically direct T cells against tumor cells (*35*). In this context, appropriate receptor expression is key for antigen recognition. With the knowledge we gained from MPRA and SONAR (Fig. 2, 3) using syn3UTRs, we tested as proof-of-concept if we could to manipulate protein expression of the CDK4-specific T cell receptor (TCR, (*36*)). The CDK4 TCR is derived from a melanoma patient, recognizing an HLA-A2 restricted neo-antigen (*36*). We first defined the optimum of SF occurrence in the syn3UTR and found that 4-6 motif occurrences drive the highest GFP expression (Fig 5B). We then fused the syn3UTR containing 6 occurrences of each of the 4 selected SFs to the CDK4 TCR, and we measured the TCR surface expression by flow cytometry in CD8^+^ T cells. As for GFP-syn3UTR constructs alike (Fig. 2F), syn3UTR containing CTTTCTT and CTCAGGT motifs consistently increased the TCR expression in CD8^+^ T cells (Fig. 5C). In contrast, TATTTA and AGAAGA motifs repressed the CDK4 TCR surface expression levels compared to scrambled syn3UTR control (Fig. 5C). Notably, the TCR mRNA expression levels of all syn3UTR constructs were equal (Fig. 5D), suggesting that the increased TCR protein expression was mediated through translational control. Indeed, aggregating the fold-change expression of mRNA and protein compared to scrambled control revealed that the CTTTCTT and CTCAGGT motifs increased the protein output, and the TATTTA and AGAAGA motifs decreased the protein output without significant changes in mRNA levels (n=3 donors, Fig. 5E).

Lastly, we tested whether the altered protein expression levels of the CDK4 TCR modulated the capacity of T cells to kill their target cells. We co-cultured melanoma cells that endogenously express the CDK4 neo-antigen (*36*) with CD8^+^ T cells expressing the indicated CDK4-syn3UTR constructs, and we measured the induction of cell death each hour for 24 h with Caspase3/7 activity dyes. Notably, CD8^+^ T Cells expressing the CDK4 TCR under control of CTTTCTT- or CTCAGGT-containing syn3UTRs were superior in killing CDK4-expressing melanoma cells compared to scrambled syn3UTR control TCR, or TCRs fused to syn3UTRs containing the AGAAGA or TATTTA motifs (Fig. 5F). In conclusion, SONAR can model and inform on the translational control of clinically relevant protein expression, which can potentiate the T cell functionality against cancer cells.

## Discussion

Protein abundance is defined by a plethora of transcriptional and post-transcriptional regulatory mechanisms (*1*). In this study, SONAR determined the importance of SFs and their contribution to protein abundance and informed the design of sequences to manipulate the amplitude, timing and cell type specificity of protein expression. We show as proof-of-concept that a TCR gene engineering approach which increased expression can also improve the T cells activity towards its target cells.

To date, efforts to optimise protein production primarily focus on codon usage and promoter strength (*37*–*40*). With SONAR models one can model how the importance of codon usage alters throughout cell types and altered cell states. We therefore envision that SONAR models could help tailor optimal protein expression in a specific cell type, and state, of interest. Notably, even in the presence of a codon-optimized GFP or CDK4 TCR combined with strong promotors (PGK and viral backbone), we could trespass the limits of protein expression, and enhance protein production solely by generating a synthetic 3’ UTR containing SFs informed by SONAR. In addition to the amplitude of expression, synthetic 3’UTRs could manipulate the timing of expression during T cell activation, an application that could safeguard expression from leaky constructs. We foresee that our methodology and custom 3’UTR designs can be successfully used in virtually all mammalian cell types and proteins of interest.

Here, we exploited our findings with SONAR to boost protein production for a clinically relevant protein, i.e. neo-antigen specific T cell receptors. We found that syn3UTRs can boost protein expression through translation control. The higher protein levels had functional consequences and improved the cytotoxic potential of T cells in response to tumor cells. We anticipate that the SONAR-guided UTR-mediated manipulation of protein production can be useful to many domains of biology.

While we provide tools to model, understand, and manipulate protein expression, we also highlight that context is key. Both cellular context (cell type, resting/activated) as well as sequence context had a drastically different effect on protein production. We believe that SONAR will greatly help in capturing the former, but more work is needed to fully capture the interplay of sequence motifs and its surrounding sequence context. A larger-scale MPRA could potentially shed light on this cross-talk of motifs and surroundings, akin to previous work on promoter motifs and surrounding sequence context of transcription factors (*17*, *37*, *41*).

We envisage that SONAR can provide important fundamental insights in many settings where the protein expression landscape is altered, such as during cell differentiation, transformation, and other stress situations such as hypoxia or starvation. This includes fundamental research questions, as recently exemplified by HLA-ligandome prediction from sequence alone (*42*). In addition, SONAR can inform the optimized design for boosting protein production for a clinically relevant protein, as exemplified here for the expression of neo-antigen specific TCR in human T cells. We foresee that SONAR-guided UTR-mediated manipulation of protein production can be applied to many domains of biology with therapeutic potential such as recombinant protein production or mRNA vaccine technologies.

## ACKNOWLEDGMENTS

We thank B. Popovic, M. Messemaker, B. Kwee, T. Schumacher for helpful discussions. We thank B. P. and N.D. Zandhuis for critical reading of this manuscript, and W. Scheper and T. Schumacher for kindly providing the CDK4 TCR construct.

## Author contribution

Conceptualization, project administration, writing: BPN, MCW. Software, data curation, analysis, visualization: BPN, KB. Validation, investigation, methodology: BPN, APJ, KB, AG, AB. Funding acquisition, supervision: MCW.

## Data and material availability

Code and scripts have been deposited on github (https://github.com/BenNicolet/SONAR-Learning-the-sequence-code-for-protein-abundance). Models were deposited on Zenodo (https://zenodo.org/record/8263086). All expression data used in this study were previously published: RNA-sequencing data retrieved from (*14*, *43*–*45*) on the National Center for Biotechnology Information (NCBI)’s gene expression omnibus (GEO) and the European Nucleotide Archive (ENA); for mass spectrometry-based proteomics and protein synthesis rate, datasets were retrieved from (*14*, *21*, *22*). Reporter assay data were deposited on GEO (https://www.ncbi.nlm.nih.gov/geo/query/acc.cgi?acc=GSE240919). See details in Table S1. All data are available in the main text or the supplementary materials.

## SUPPLEMENTARY MATERIALS

Materials and Methods

Supplementary Text

Figs. S1 to S8

Tables S1

References (1-24)

Data S1 to S6

