## Supplementary material for "Learning the sequence code of protein expression in human immune cells": Fig S

### Supplementary Materials for:
### Learning the sequence code for protein abundance in human immune cells

Benoît P. Nicolet*, Anouk P. Jurgens, Kaspar Bresser, Aurélie Guislain, Antonia Bradariç, and Monika C. Wolkers*

**This PDF file includes:**

Materials and Methods
Supplementary Note
Figure S1 to S8 and captions
Table S1
Captions for Data S1 to S6

**Other Supplementary Materials for this manuscript include the following:**

**Data S1 to S6:**

Data S1. SONAR feature importance from protein abundance models

Data S2. Cell-type specific SF usage of protein abundance models

Data S3. Syn3UTR sequences

Data S4. Massively parallel reporter assay motif enrichment in HEK293, CD4+ and CD8+ T cells

Data S5. SONAR feature importance from mRNA abundance, translational rate, protein abundance in CD4+ T cells throughout activation

Data S6. Relation of mRNA abundance and putative m7G sites in HeLa cells

Materials and Methods

**Data acquisition.** For analysis of mRNA abundance in HEK293T, HeLa and K562 cell lines, and in primary peripheral blood-derived immune cells, we used RNA-sequencing data retrieved from (*1–4*) on the National Center for Biotechnology Information (NCBI)'s gene expression omnibus (GEO) and the European Nucleotide Archive (ENA), see details on datasets in Table S1. For Mass spectrometry-based proteomics and protein synthesis rate, datasets were retrieved from (*3*, *5*, *6*) and acquired from the respective supplementary materials (details in Table S1).

**RNA-seq and mass spectrometry analysis.** For RNA-seq analysis, samples were quasi-mapped using Salmon ((*7*) version 1.4) on the human coding transcriptome hg38-release 92 from ENSEMBL (*8*). The normalized counts in 'Transcripts per kilobase per million' (TPM; which corrects for transcript length and library size) was calculated by Salmon, zero-count transcripts were removed, and ENSEMBL annotations (r92) were used to filter for protein-coding transcripts. TPMs were log10-transformed, and biological replicates were used separately as input for modelling. Of note, replicate 2 of K562 RNA-seq sample was removed due to poor correlation with other K562 samples (~0.7 versus >0.9 Pearson's correlation coefficient for other biological replicates). For mass spectrometry analysis, protein copy numbers (CN) were used as a measure of protein abundance. CN values were retrieved from supplemental information of indicated publications and log10 transformed. When the pairing information of RNA-seq / proteomics replicates was not available, we averaged TPM and CN per cell type or activation status (n=3-4 biological replicates) in Fig 1,4,5.

**SONAR pipeline:**

**A) Feature library construction.** The sequence feature library was built using the 94,348 protein-coding mRNA isoform sequences acquired from ENSEMBL (release 104, accessed August 2021; see details in Fig. S1A). To annotate the separate occurrence of sequence features in the coding region (CDS) and the 5' and 3' Untranslated Regions (UTR), we acquired the raw sequence for each of these regions from ENSEMBL Biomart (*9*). For each mRNA transcript, we searched for the following sequence features: **A)** For the full mRNA, we included sequence homology between humans and zebrafish (*Danio rerio*, acquired from ENSEMBL BiomaRt) and miRDB miRNA seed scores (*10*) of miRNAs that were detected in T cells (*11*). **B)** For the CDS, we extracted the codon and amino acid usage using coRdon (*12*), expressed as a percentage occurrence for the CDS. **C)** For each mRNA region (5'UTR, CDS, 3'UTR) separately, we searched for the percentage of GC content, mRNA length (in nucleotides), and for putative mRNA modification site occurrence (m1A, m5C, m6A, m7G) from the RMVar database (accessed at https://rmvar.renlab.org/ in June 2021. We also included RNA-binding protein (RBP) binding motifs from the 142 RBPs annotated in the ATtracT database (*13*) for each mRNA region separately. All features combined resulted in a library of 7112 SFs (Fig S1A).

Furthermore, we included 13 putative post-translational modification (PTM) sites into the models from the *dbPTM* database (only PTM with >1000 annotated observations were considered; accessed at https://awi.cuhk.edu.cn/dbPTM/ in June 2021 (*14*)). This entailed, amongst others, Glycosylation, Palmitoylation, Phosphorylation, S-nitrosylation, Succinylation, Sumoylation, Ubiquiniation site occurrences (Fig S1A). For mRNA abundance modelling, the putative PTM sites were omitted. Of note, because more than one mRNA isoform can lead to a similar protein isoform, the feature library was averaged per UniProt identifier to allow for protein abundance modelling. Furthermore, post-transcriptional mRNA modification sites (e.g. m6A, …) and protein PTM inform on the occurrence of a putative modification site in the mRNA or protein. They are not a measurement of the actual modification of the indicated site, nor do they include information on the percentage of mRNA or protein modification that occur.

**B) Model selection, optimization, training, and testing.** To prevent data leakage, training (80%) and testing (20%) sets were split at random prior to modelling. XGBoost (*15*) ran in Caret (*16*) in R where hyperparameters were tuned in a grid fashion by varying: *'nrounds'* from 1 to 10000 (best: 10000); '*max_depth'* from 1 to 300 (best: 6); *'colsample_bytree'* from 0 to 1 by steps of 0.1 (best: 0.4); *'eta'* from 0 to 0.5 by steps of 0.05 (best: 0.05); *'gamma'* from 0 to 10 in steps of 1 (best: 1); *'min_child_weight'* from 0 to 1 by steps of 0.1 (best: 0.9); *'subsample'* from from 0 to 1 by steps of 0.1 (best: 1). Of note, *gamma* which controls for the loss reduction at the node level (the higher the *gamma* parameter, the more conservative the model), was kept at 0 for models depicted in Fig 4&5 to allow for 'deeper' models (which include information on all sequence features in the final feature importance computation whether or not these pass the gamma threshold). In turn, this allowed for a more complete comparison of feature importance between models in Fig 5. A 5-fold cross-validation (CV) was used to test the models' robustness. Hyperparameter tuning was performed on the training set (i.e. $CD8^+$ Tn) only, and the target metric ('Rsquared') was assessed on the held-out test set within the 5-fold CV (training set only). Please see our GitHub repository for details on the parameters, and to Zenodo for the models.

**C) Downstream analysis of SONAR models.** For each model, the final model performance was determined using the held-out test set (R squared). Feature importance was extracted using standard Caret functions (*'varImp()'*; no scaling was applied; *'scale=FALSE'*), and $log_{10}$-transformed. To allow for better appreciation of feature importance scores, the minimum value in the datasets was subtracted to the feature importance, such that feature with lowest importance becomes the value '$10^0$' (=1), and that all values ended up in the positive number space. Uniform Manifold Approximation and Projection (UMAP) was obtained with *'umap'* package (T. Konopka, version 0.2.9.0). For comparison of the relative feature importance between mRNA, translation rate and protein models (Figure 5), $Log_2$ fold-change (LFC) was computed as LFC(AB) = LFC(B) - LFC(A). K-means clustering was performed using pHeatmap (*17*) version 1.0.10. For information on feature grouping based on variation, please see supplemental note #1 below.

**Analysis of m7G sites in HeLa cells.** To compare the percentage of m7G sites to mRNA abundance in HeLa cells (*4*), we re-analysed the G to T/C misincorporation data set from *Zhang et al.* (*18*). We also calculated the occurrence of m7G sites in the 655 mRNAs that were identified in (*18*). Correlation and statistics were computed using *'cor.test()'* of the R 'Stats' package.

**Isolation of cell type-specific sequence features.** To isolate the cell type-specific SFs of protein models, we used XGBoost with adapted hyperparameter to separate cell types. Subsequently, the feature importance was extracted. Of note, in this case the feature importance is a measure of the SF contribution to tease apart cell types. SFs were then ranked by contribution, and the top 500 most contributing SFs were used for visualization. For details, please refer to scripts on GitHub.

**Generation of synthetic 3'UTRs.** To generate 3'UTRs containing SFs identified with SONAR, we first generated a scrambled sequence backbone of 180nt that was devoid of important sequence features, by iteratively (3 rounds) searching for motifs predicted by SONAR to have high feature importance in protein models. Identified motifs were mutated by modifying 2 nucleotides. To test the effect of sequence motifs on protein expression, 6 occurrences of 6- and 7-mer motifs predicted to have high importance (467 motifs) were inserted into the scrambled sequence by replacing internal nucleotides to keep sequence length identical (180nt). For measuring the cell-type specificity of motifs and activation induced changes in GFP, gene blocks were generated (IDT technologies) with the following 4 motifs (CTCAGGT (7-mer), CTTTCTT (7-mer), AGAAGA (6-mer), TATTTA (6-mer)), inserted into the scrambled 3'UTR or for CXCL2 (138-204nt)

WT and padded on the 5' and 3' end with restriction sites described below. Sequences of oligos are listed in Data S3.

**Cloning.** For the library and single motif testing, 3′UTR sequences were cloned into NotI and BamHI restriction sites of pRETRO-SUPER_GFP (*19*) at the 3' end of the GFP CDS, as described before (*20*). Plasmids were amplified in DH5a high competency (NEB Cat #C2987H) according to the manufacturer's protocol, and DNA was isolated with NucleoSpin Plasmid Transfection-grade DNA extraction kit (Macherey-Nagel Cat #740490.250). To generate HLA-A*02:01-restricted $CDK4_{R24L}$ T cell receptor containing synthetic 3'UTRs, the BamHI restriction site was used downstream of the stop codon of the TCR in the pMP71 vector containing the CDK4 TCR (*21*). All sequences were confirmed by Sanger sequencing.

**Virus production.** FLYRD18 retroviral packaging cells (ECACC 95091902) were cultured in culture-supplemented IMDM (Gibco-BRL), containing 10% fetal bovine serum (FBS; Bodinco), 2 mM L-glutamine, 20 IU/mL penicillin G sodium salts, 20 µg/mL streptomycin sulfate (all Sigma Aldrich) in a humidified incubator at 37°C + 5% $CO_2$. Cells were pre-plated at a density of $1 \times 10^5$ per well in a 6-well plate for 16 h and then transfected with indicated plasmid using GeneJammer (Agilent Cat #204130) according to manufacturer's protocol. Retroviral supernatant was harvested after 48 h, and was either used freshly, or was snap-frozen in liquid nitrogen and stored at -80 ºC for later use.

**Cell culture.** Studies with human T cells from anonymized healthy donors were performed in accordance with the Declaration of Helsinki (Seventh Revision, 2013) after written informed consent (Sanquin). Peripheral blood mononuclear cells (PBMCs) were isolated through Lymphoprep density gradient separation (Stemcell Technologies) and cryopreserved until further use. Human T cells, and K562 cells (ECACC 89121407), were cultured in complete IMDM medium (IMDM (Gibco-BRL), supplemented 10% fetal bovine serum (FBS; Bodinco), 2 mM L-glutamine, 20 IU/mL penicillin G sodium salts, 20 µg/mL streptomycin sulfate (all Sigma Aldrich)). HeLa (ECACC 93021013), and HEK293 (ECACC 85120602) cells were cultured in DMEM (Gibco-BRL), containing 10% fetal bovine serum (FBS; Bodinco), 2 mM L-glutamine, 20 IU/mL penicillin G sodium salts, 20 µg/mL streptomycin sulfate (all Sigma Aldrich). Retrovirally transduced human T cells were maintained at a density of $0.5\text{-}1 \times 10^6$ cells/mL in culture-supplemented IMDM containing 100 IU/mL recombinant human (rh) IL-2 (Proleukin, Novartis), and 10 ng/mL rhIL-15 (Peprotech). Medium was refreshed every 2 days. Cell lines were split every second day to maintain a confluency of max 80%.

**IncuCyte Killing Assay**. Human $CD8^+$ T cells were retrovirally transduced with the low affinity T cell receptor (TCR) recognizing the CDK4R>L neoantigen (clone 55) containing syn3UTRs (*21*). $CD8^+$ T cells were FACS sorted based on TCR expression (mTRBC monoclonal antibody -PE-Cy7 (clone H57-597; eBioscience #25-5961-82)). FACS-sorted $CD8^+$ T cells expressing the $CDK4_{R>L}$ TCR were rested overnight and then co-cultured for 24 hours with Mel526 tumor cells naturally expressing the $CDK4_{R>L}$ neoantigen, in a 1:1 ratio. 5µM Caspase-3/7 green apoptosis assay reagent (Sartorius #4440) was added at the start of the experiment in complete IMDM medium to the Mel526 cells 30min before T cells were added, as described by manufacturer. As a negative control, tumor cells were cultured without T cells. One picture was taken every hour by the IncuCyte Live-Cell analysis System (Sartorius) for a total of 24h. T cell killing was assessed with the integrated IncuCyte Base Analysis Software (Sartorius) using dead tumor cell count. Importantly, each data timepoint was normalized to the negative control to account for tumor death without T cells.

**Retroviral transduction**. Human PBMCs were thawed, washed with culture medium, and rested for 1 h at 37 ºC. T cells were activated for 48 h with αCD3/ αCD28 as previously described (*22*). Briefly, 24-well plates were pre-coated overnight with 0.5 µg/mL rat α-mouse IgG2a (clone MW1483; Sanquin) at 4 °C, washed once with PBS, coated with 1 µg/mL αCD3 (clone Hit3a, eBioscience) for a minimum of 3 h at 37 °C, and washed once with PBS. $1.3 \times 10^6$ PBMCs/well were plated in IMDM medium supplemented with 1 µg/mL αCD28 (clone CD28.2; eBioscience) for 48 h in a humidified incubator at 37ºC + 5% $CO_2$. Cells were collected, and retroviral transduction was performed as previously described (*23*). Briefly, non-tissue cultured treated 24-well plates were coated overnight with 50µg/mL Retronectin (Takara) at 4°C, and washed once with PBS. 400µL viral supernatant was added per well, and plates were centrifuged for 45 min at 4 °C at 4500 rpm (2820g). 1 x $10^6$ activated T cells, or K652 cells were added per well, plates were centrifuged for 5 min at 1000 rpm, and incubated overnight at 37 °C. HeLa and HEK293 cells were plated at a confluency of 2.5 x $10^5$ per well in a 6-well plate and cultured overnight at 37 °C + 5% $CO_2$. The next morning, prior to retroviral transduction with 400µL virus supplemented with 10 µg/mL polybrene (Merck Millipore). Plates were centrifuged for 5 min at 1000 rpm and incubated overnight at 37 °C. The next day, culture medium of all cells was refreshed, and cell culture was performed as described above.

**Flow Cytometry.** Four days post retroviral transduction, or after 6 or 16h activation as described above (for T cells only), cells were harvested and washed with FACS buffer (PBS, 1% FBS, 2 mM EDTA). Cell lines were analyzed for GFP expression. T cells were labeled with α-CD4 (clone SK3), α-CD8 (clone SK1; all Biolegend) and analyzed for GFP expression. The expression of the $CDK4_{R24L}$ TCR was determined with mTCRβ monoclonal antibody (clone H57-597) labeled with PE-Cy7 (eBioscience Cat #25-5961-82) as previously described (*23*). Near-IR live/dead marker (Life Technologies) was used to exclude dead cells from analysis. Cells were acquired on a Symphony flow cytometer (BD Biosciences); data were analyzed using FlowJo (FlowJo LLC, version 10).

**3'UTR library cloning.** A library of 467 synthetic 3'UTRs, each containing 6 repeats of a distinct sequence motif, was synthesized as an oligo pool by Twist Bioscience. Each oligo was designed as outlined in Figure 4A and was flanked by a 5' BamHI site and a 3' NotI site (Data S7). To generate a dsDNA library 10 ng of the oligo pool was PCR amplified for 9 cycles with NEBnext polymerase (NEB) using primers (Forward: 5'-TGGCATTGGATCCCTATTCC-3', Reverse: 5'-TAATAAGCGGCCGCGATGAG-3') adding short overhangs to facilitate restriction digest. Resulting product and the pRETRO-SUPER plasmid were digested using BamHI-HF (20U, NEB) and NotI-HF (20U, NEB) at 37°C for 2 h. During the last hour of the pRETRO-SUPER digest, 5U Quick CIP (NEB) was added to dephosphorylate the backbone. The library insert and the backbone were isolated and purified from a 1.5% agarose gel, and subsequently ligated at a backbone:insert ratio of 1:2 using 40U T4 ligase (NEB) using a cycling ligation protocol (20 cycles of: 5 min at 4°C, 10 min at 26°C, 5 min at 4°C and 40 min at 16°C). Ligation product was amplified through chemical transformation into 5-alpha Competent E. coli (NEB) and subsequent culture at 32°C. The performed UTR library cloning maintained a 123x coverage, with an approximate 0.67% contamination of re-ligated plasmid backbone. Presence of all synthetic 3'UTRs was confirmed through deep-sequencing.

**3'UTR parallel reporter assay.** Retroviral library was produced by transfecting 4.5 μg of the library plasmid DNA into FLYRD18 retroviral packaging cells, as described above. HEK and HeLa cells were cultured to a density of 60% prior to addition of virus-containing supernatants, and subsequently expanded for 10 days. PBMCs were activated with αCD3/ αCD28 and spinfected with virus-containing supernatants, as described above, and expanded for 10 days in RPMI supplemented with 10% fetal bovine serum (FBS; Bodinco), 2 mM L-glutamine, 20 IU/mL penicillin G sodium salts, 20 μg/mL streptomycin sulfate (all Sigma Aldrich), Non-Essential Amino Acids (Gibco), Sodium pyruvate (Gibco), 50 IU/mL recombinant human (rh) IL-2 (Proleukin, Novartis), 5 ng/mL rhIL-15 (Peprotech), and 10 ng/mL rhIL7 (Miltenyi). Transduction efficiencies were kept between 5-10% to minimize multiple transgene integrations. After expansion, cells were harvested and stained with Near-IR live/dead marker (Life Technologies) for dead cell exclusion, in addition to anti-CD8-AF700 (Biolegend) and anti-CD4-BV510 (Biolegend) in case of the T cell assay. The top and bottom 15% GFP expressing cells, in addition to the full GFP-positive fraction, were sorted on an Aria II (BD Biosciences). FACS-sorted cells were subsequently pelleted and resuspended in 1 µl DirectPCR Lysis Reagent (Viagen) per 2,500 cells, supplemented with 0.5 mg/ml proteinase K (Viagen). The obtained lysate was incubated at 55 °C for 3 h, followed by 85 °C for 45 min, and 95 °C for 5 min.

**3'UTR reporter assay library amplification and sequencing.** An initial pre-amplification of the synthetic 3'UTRs was performed on 20 µl lysate (±50,000 cells) in a 100 µl reaction volume using the NEBnext (NEB) and 10 µM forward (5'-AAAGACCCCAACGAGAAGC-3') and reverse (5'-AGTCTATAGCTACTAGGCG-3') primers. An initial touchdown stage was performed dropping the annealing temperature 1°C per cycle from 58°C to 50°C (8 cycles), followed by 12 cycles at 50°C. Extension time was kept at 20 seconds. This initial PCR was performed in duplicate for each sample, as a technical control. Excess primers were removed from each reaction by incubating 20 µl of the initial PCR volume with 40 U Exonuclease I (NEB) at 37°C for 15 min, followed by a heat-inactivation at 80°C for 15 min. Each of these samples were subsequently used as template for a second PCR, with primers (Forward: 5'-ACACTCTTTCCCTACACGACGCTCTTCCGATCTAGACCCCAACGAGAAGCGCGATCAC-3', Reverse: 5'-GACTGGAGTTCAGACGTGTGCTCTTCCGATCTAGTCTATAGCTACTAGGCGATA-3') designed to amplify the product and attach partial Illumina adapters. This second PCR was performed using the NEBnext (NEB) and consisted of 16 cycles with a constant annealing temperature of 65 °C and extension time of 30 sec. Correct products (±275 base pairs) were purified from a 2% agarose gel using the Monarch DNA Gel Extraction Kit (NEB). Purified products were sequenced using the Amplicon-EZ service from Genewiz (Azenta Life Sciences; PE 250 cycles) or the genomics core facility of the Netherlands Cancer institute (PE 150 cycles). Raw data are available on GEO (see below & Table S1).

**3'UTR parallel reporter analysis.** Sequencing adapters were trimmed using cutadapt using the settings -n 2, -e 0.2 and -M 62. Resulting sequences were aligned to the UTR library using bowtie2 in local mode. PCR hybrids that were generated during library amplification were removed from the alignment file using a custom script. Read counts were obtained using htseq-count. Finally, sequence motifs detected with less than 10 reads across all samples were excluded from the analysis, and individual libraries were normalized to 10,000 reads. HeLa samples displayed a poor replicate correlation and were excluded from our analysis (data are available on GEO).

**Statistical analysis and Data visualization.** Statistical analysis was performed in R or with GraphPad PRISM using the test indicated in the respective Figure legends. Differences were considered significant if adjusted p-value <0.05, and are depicted in each graph. Plots were generated with ggplot2 (*24*) version 3.0, and with GraphPad PRISM version 9. Heatmaps were generated with pHeatmap.

**Supplemental text:**

**Note #1:** For data analysis, we used 'corrected S.D.' in Fig. S2, S6, for the following reason. To determine how feature importance varies between the models of different cell types (or time points for T cell activation), we required a measure of variation for each feature that could be extracted and compared to other features. We first calculated the standard deviation (S.D.), which measures the distance from the average values. However, because feature importance had a broad range of values (from 0 to ~$10^{10}$ (arbitrary units)), the S.D. of highly important features was masking the ones of lesser importance. As an example, a feature with an average feature importance of 100 and an S.D. of ± 5 will have the same S.D. as a feature with an average importance of 10 ±5. Yet, the variation of the former is 5%, while this of the latter is 50%. To correct for this bias introduced by the broad range of feature importance, we divided the S.D. of features by their respective average (throughout the models of different cell types). This resulted in a 'corrected S.D.' (eq. 1 below), which was immune to the actual values of feature importance, yet which retained the degree of variation per features.

Equation 1:

$$corrected\ S.D. = \frac{\sigma_{feature}}{\mu_{feature}}$$

Where $\boldsymbol{\sigma}$ is the S.D. and $\boldsymbol{\mu}$ is the average feature importance (computed per feature).

**Fig.S1 Nicolet et al.**

**A**

| features | 5'UTR | CDS | 3'UTR | full mRNA | full protein | total mRNA | total protein |
|---|---|---|---|---|---|---|---|
| **RBP motifs** (142 RBP; Attract) | 2247 | 2270 | 2266 | 0 | 0 | 6783 | 6783 |
| **GC %** | 1 | 1 | 1 | 0 | 0 | 3 | 3 |
| **length** | 1 | 1 | 1 | 0 | 0 | 3 | 3 |
| **miRNA seed score** (miRDB) | 0 | 0 | 0 | 218 | 0 | 218 | 218 |
| **mRNA modification/editing predicted sites** (m1A, m5C, m6A, m7G, A-to-I, from RmVar) | 5 | 5 | 5 | 5 | 0 | 20 | 20 |
| **Sequence homology to D.rerio** | 0 | 0 | 0 | 1 | 0 | 1 | 1 |
| **codon usage** (per codon) | 0 | 64 | 0 | 0 | 0 | 64 | 64 |
| **codon usage** (per encoded amino acid) | 0 | 20 | 0 | 0 | 0 | 20 | 20 |
| **Post-translational modifications** (Acetylation, Amidation, Hydroxylation, Malonylation, Methylation, N_linked_Glycosylation, O_linked_Glycosylation, Palmitoylation, Phosphorylation, S_nitrosylation, Succinylation, Sumoylation, Ubiquiniation) | 0 | 0 | 0 | 0 | 13 | 0 | 13 |
| **total** | 2254 | 2361 | 2273 | 224 | 13 | 7112 | 7125 |

**B**

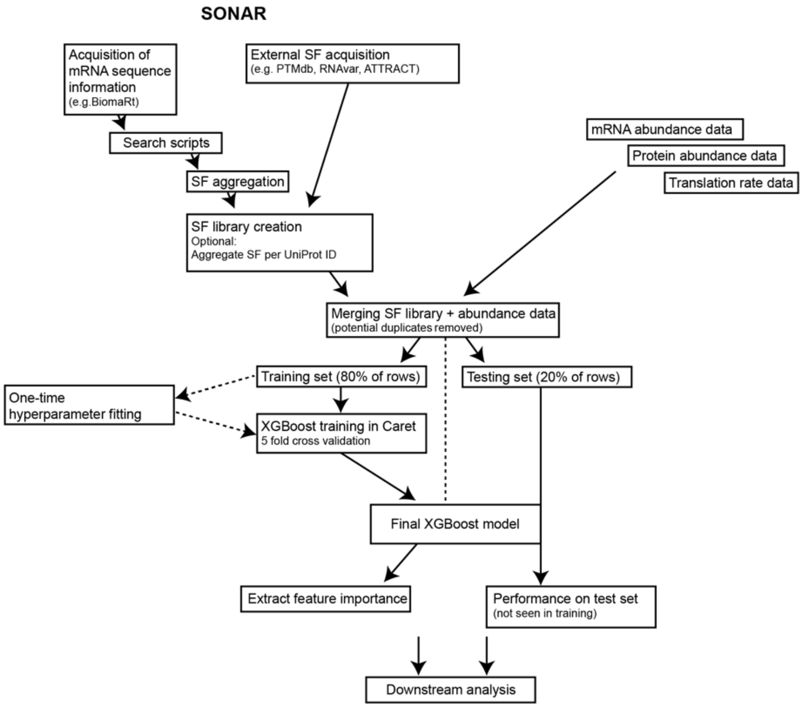

**C**

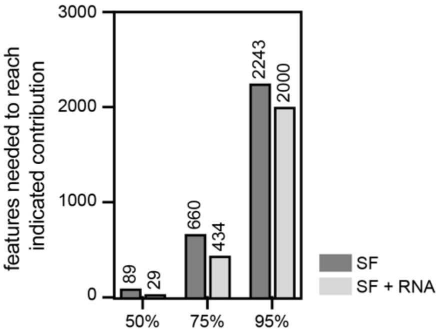

**Fig. S1: SONAR predicts protein abundance based on sequence features.**

(**A**) Table depicts the number of sequence features (SFs) included from each category in the SF library used for mRNA and protein modeling, resulting in a total of 7112 and 7125 SFs for mRNA and protein abundance prediction, respectively. (**B**) Diagram depicting the main steps in SONAR modelling. (**C**) Number of SFs needed to reach 50%, 75% and 95% cumulative contribution level to protein (SF) and SF models supplemented with mRNA abundance measurements (SF+RNA) models for all tested immune cells and cell lines combined.

**Fig. S2 Nicolet et al.**

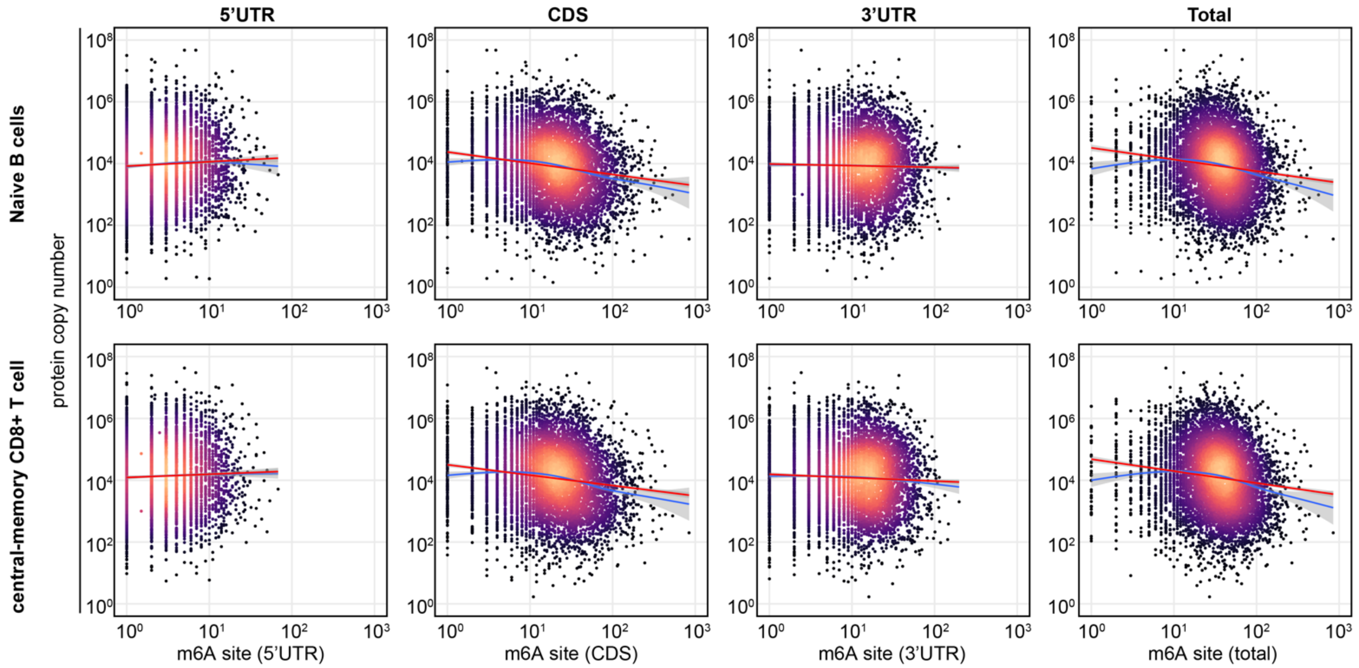

**Fig. S2: Association of protein abundance and putative m6A sites in human lymphocytes.**

Dot plots showing the relation between m6A mRNA modification site occurrence in 5'UTR, CDS, and 3'UTR or in the whole mRNA (total) and the protein abundance (copy number) in naïve B cells and in central memory $CD8^{+}$ T cells (population chosen at random). Red line depicts the linear regression, blue line depicts the generalized additive model. CN data are the average of 4 biological replicates.

#### Fig. S3 Nicolet et al.

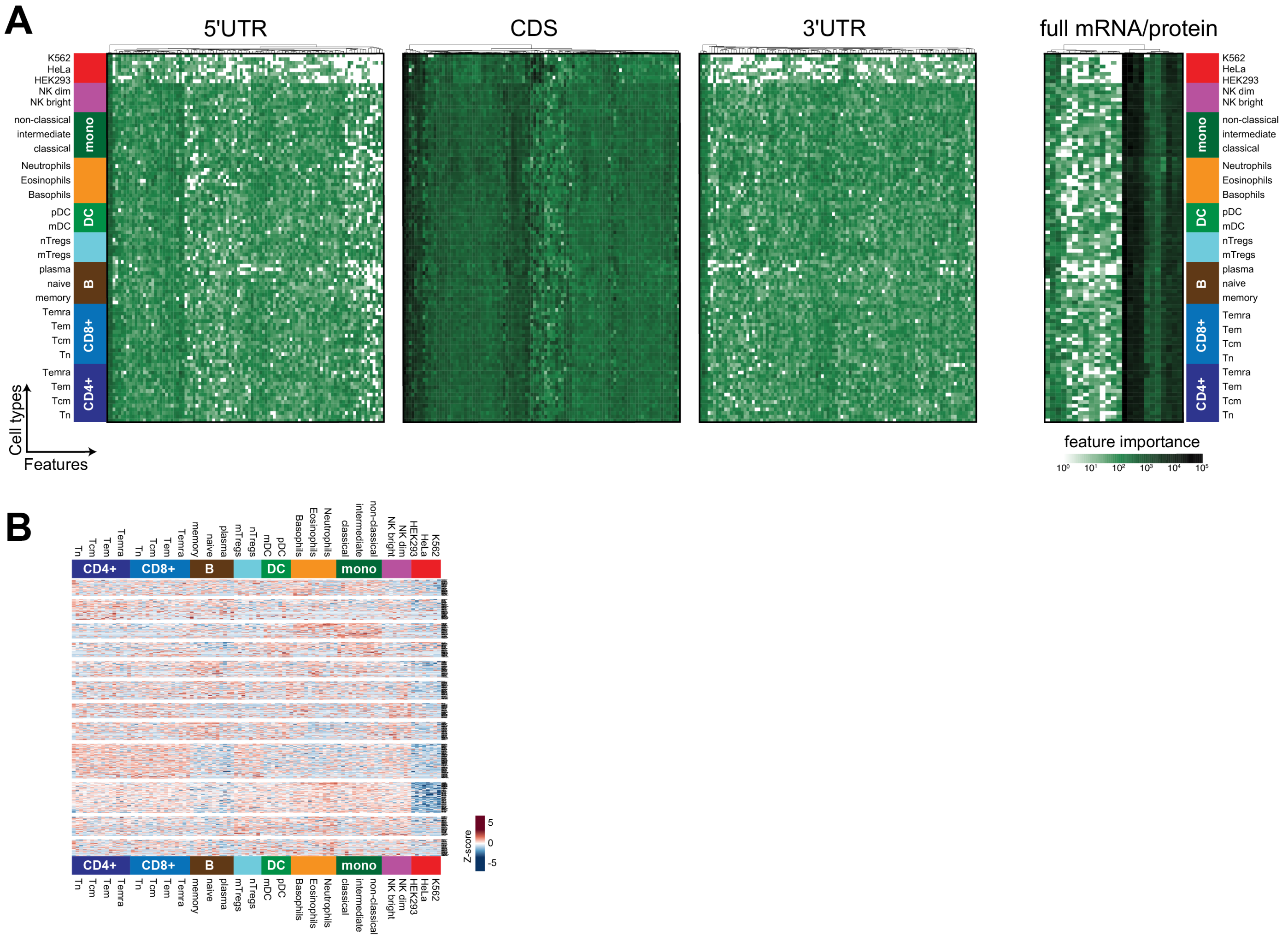

**Fig. S3. Cell type-specific SFs importance of sequence features uncovered by SONAR protein abundance models.**

(**A**) The top 100 contributing features to protein abundance of models in Fig 1 are show per mRNA region (Top 25 for 'full mRNA/protein'). (**B**) Heatmap showing the top 500 most contributing SFs used to distinguish cell types-specific SFs (XGB classifier, see methods). Color indicate SFs Z-score (per row).

Fig. S4 Nicolet et al.

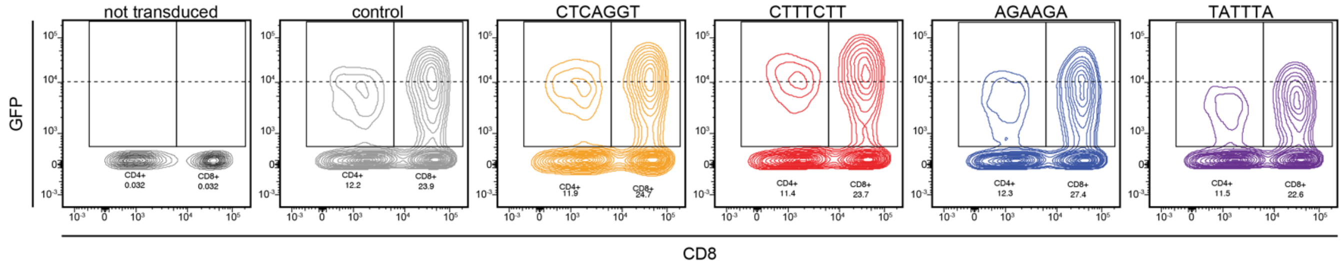

**Fig. S4. Manipulation of GFP protein expression with 3’UTR (syn3UTR) motifs.**

Human T cells were retrovirally transduced with 3’UTR reporter GFP constructs (see methods) containing 6 repeats of the indicated motifs within the 3’UTR, or with the scramble 3’UTR control. GFP expression was measured by flow cytometry. Individual panels depict CD8$^{+}$ T cells on the right, and CD4$^{+}$ T cells (depicted as CD8$^{-}$) of the same donor on the left. Numbers indicate percentage of GFP expressing cells within each T cell population. Dot plots are representative of 5 donors.

Fig. S5 Nicolet et al.

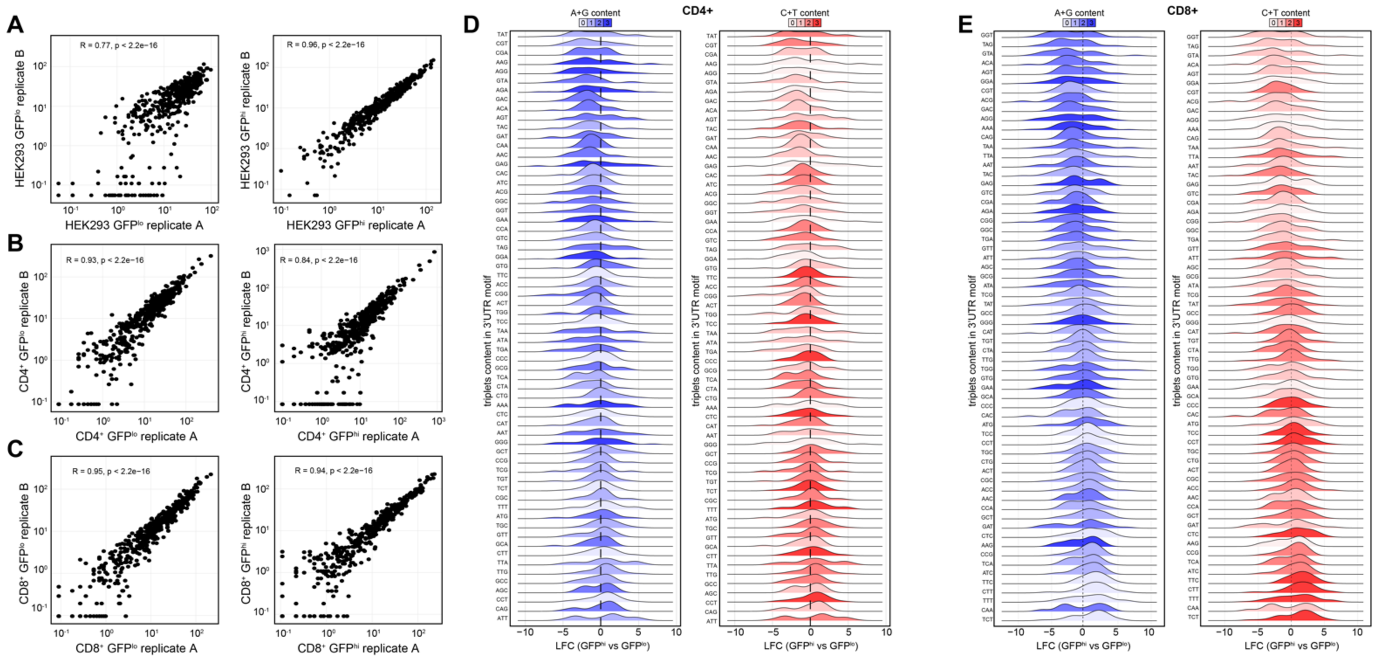

**Fig. S5. 3'UTR massively parallel reporter assay in HEK293, $CD4^+$ and $CD8^+$ T cells.**

(**A-C**) Synthetic 3'UTR (syn3UTR) massively parallel reporter assay (MPRA). Syn3UTRs containing 6 occurrences of the same 6- or 7-mer motif (n=467) were fused to the 3'end of a GFP reporter gene. The resulting syn3UTR library was introduced into (A) HEK293, (B) $CD4^+$ T cells or (C) $CD8^+$ T cells. Cells were sorted based on GFP expression (top 15%: $GFP^{hi}$ or bottom 15%: $GFP^{lo}$). Samples were split post DNA extraction as technical replicates (replicate A or B). (A-C) Shows the read count for technical replicates in all indicated cell types. Pearson's correlation coefficient and adjusted p-value are indicated. (**D-E**) Log2 fold-change (LFC) enrichment of motifs found in $GFP^{hi}$ or $GFP^{low}$ in (D) $CD4^+$ and (E) $CD8^+$ T cells. Color coding in (D-E) indicate the C+T or A+G nucleotide content within motifs.

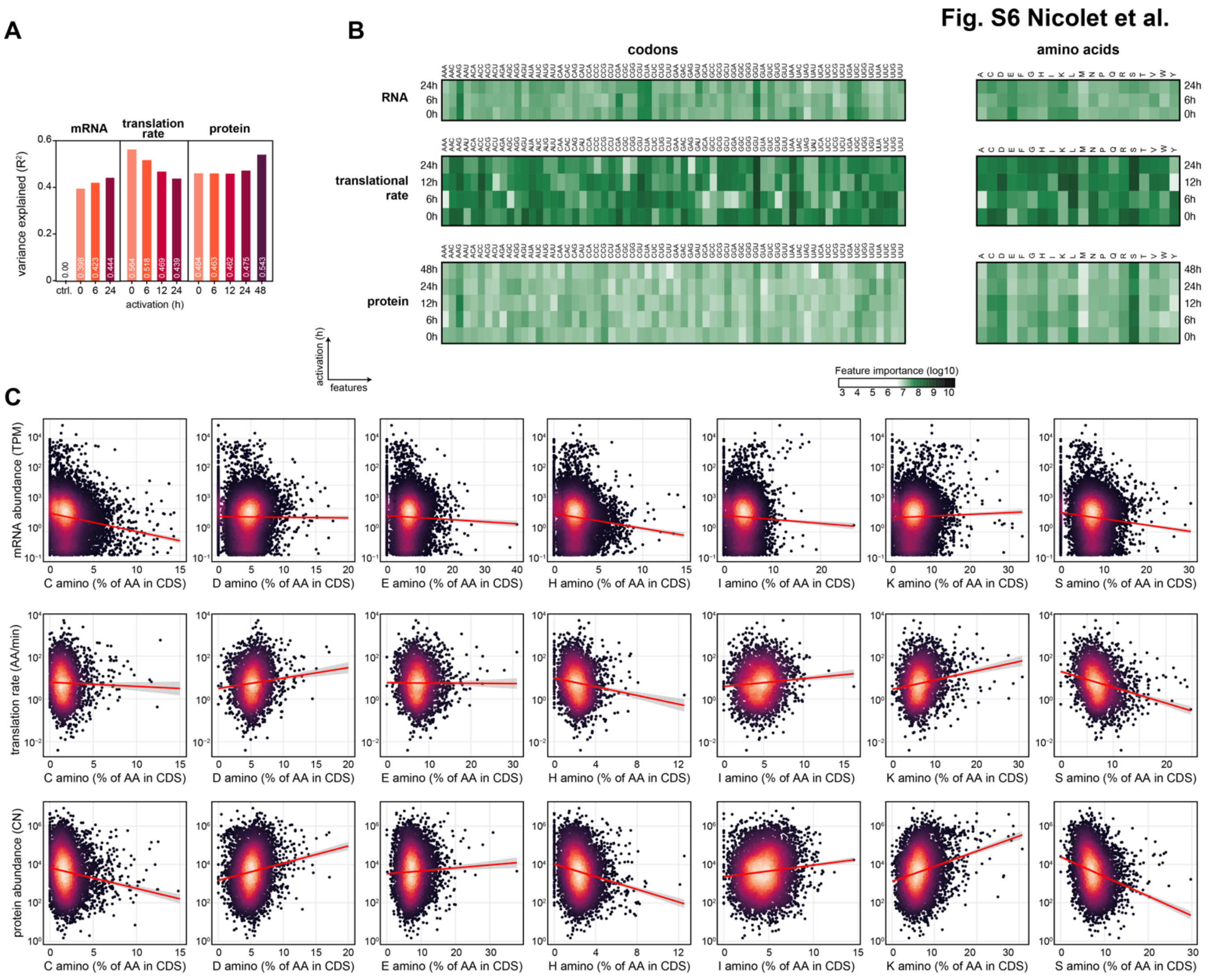

**Fig. S6. SONAR reveals activation-dependent contribution of codons and amino acids.**

(**A**) SONAR models of mRNA abundance, translation rate, and protein abundance data of CD4$^+$ T cell during activation ((*3*); average of 3 donors across timepoint). (**B**) The contribution of codon and amino acid usage on naïve human CD4$^+$ T cells upon activation was extracted from models in (A) represented in heatmaps for mRNA abundance, translation rate and protein abundance. (**C**) mRNA abundance (top panels; in Transcript per kilobase per million; TPM), translation rates (middle panels; amino acid per minute; AA/min) and protein abundance (bottom panels; in copy number; CN) are plotted against the occurrence of indicated amino acids in CDS. Abundance data are averaged across timepoints. Red line represents the linear regression of the data.

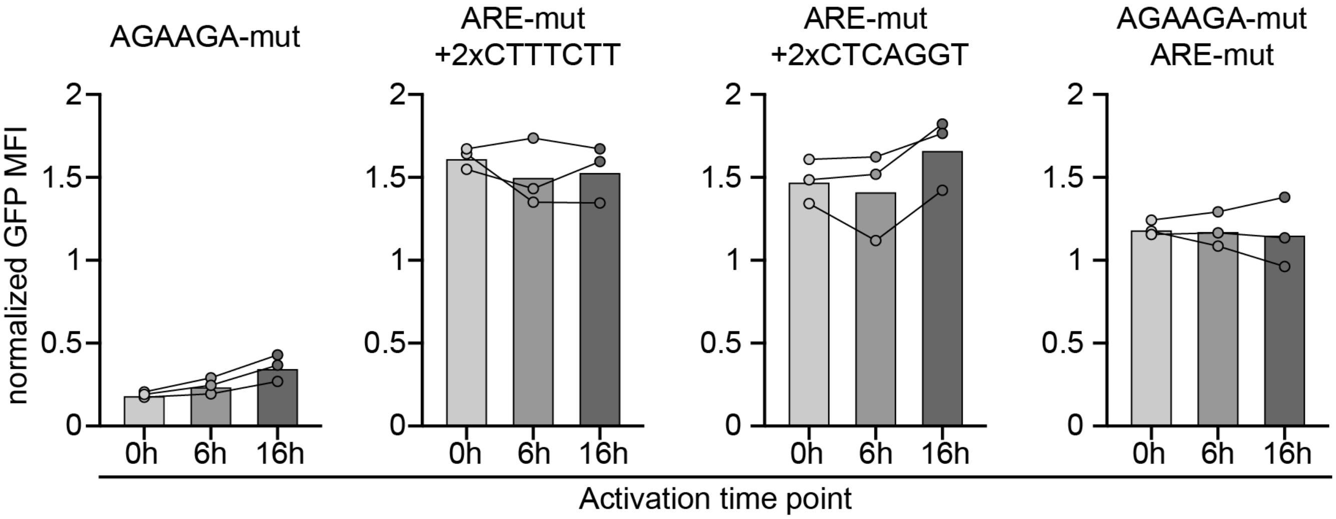

**Fig. S7. GFP expression under the control of CXCL2 3'UTR mutants through T cell activation.**

Human T cells were retrovirally transduced with CXCL2 3'UTR mutants GFP reporter constructs (*see methods*) where the natural AGAAGA element was mutated alone ($AGAAGA_{mut}$), or together with the ARE ($AGAAGA_{mut}$ $ARE_{mut}$). In addition, the ARE mutant ($ARE_{mut}$) was reconstituted with two CTTTCTT or CTCAGGT motifs were reintroduced into the ARE locus while keeping the length intact ($ARE_{mut}$ + 2x CTTTCTT or $ARE_{mut}$ + 2x CTCAGGT). GFP expression was measured by flow cytometry at 0h (rest) or after T cell activation for 6h or 16h. GFP was normalized to the scramble 3'UTR control of the corresponding timepoint.

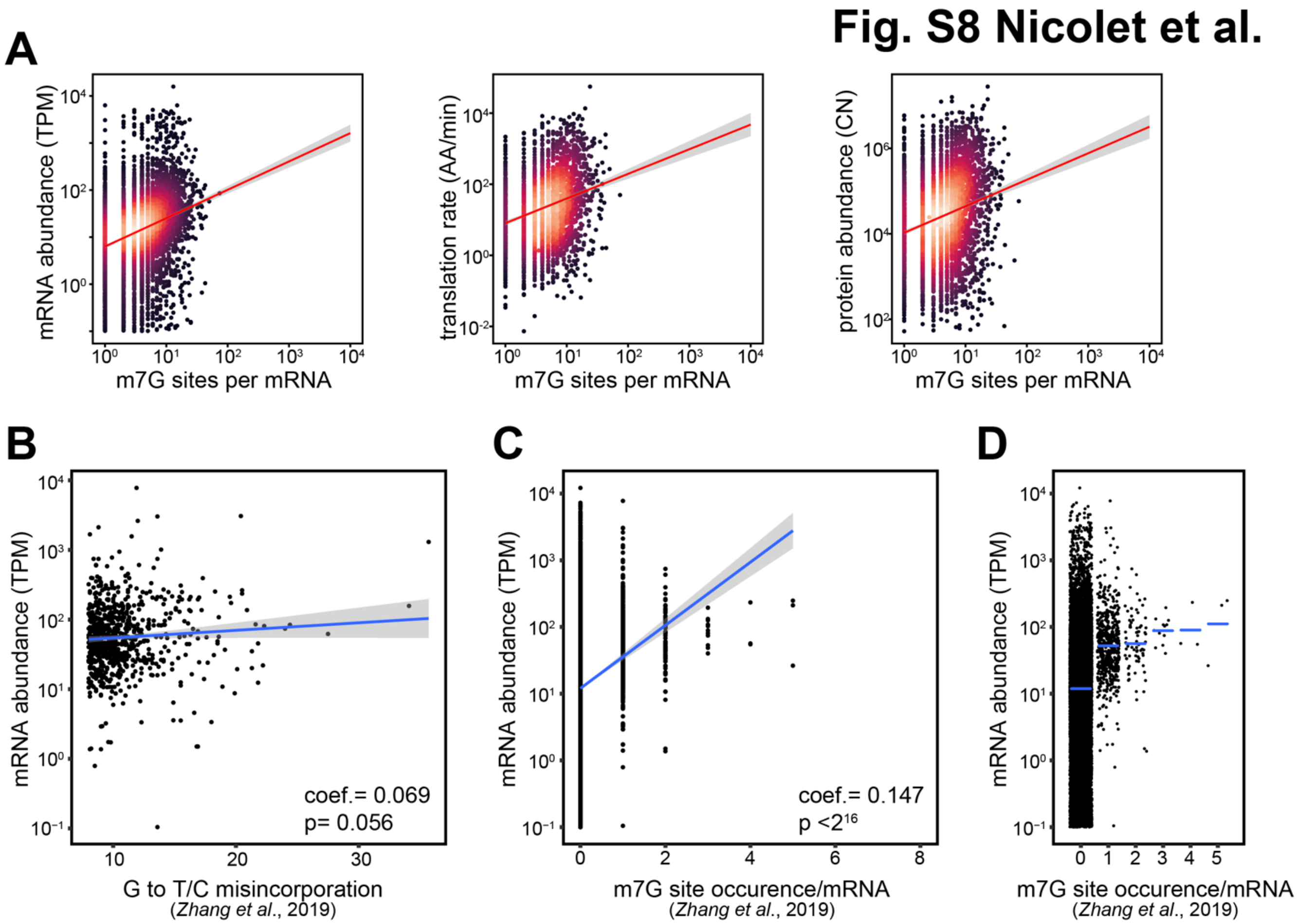

**Fig. S8. m7G site occurrence, but not modification status is correlated with mRNA abundance in HeLa cells.**

(**A**) Relation between mRNA abundance, translation rate and protein abundance with the occurrence of putative m7G sites per mRNA. (**B**) m7G-mediated misincorporation data from Zhang et al., (***18***) were re-analyzed and plotted against mRNA abundance data of HeLa cells from Martinez et al. (*4*). (**C-D**) mRNA abundance versus concatenated data of Zhang et al., into m7G site occurrence per mRNA in (**C**) a scatter plot and (**D**) a grouped dot plot. Blue lines in (B) and (C) depict the Pearson's correlation for which the coefficient and the adjusted p-value are indicated. Blue bar in (D) depicts the mean per group.

**Table S1. List of data source**

| used in | cell type | type of data | reference | accession no. |
|---|---|---|---|---|
| Fig. 1 | Immune cells | RNA-seq | Monaco et al., 2019 | GSE107011 (GEO) |
| Fig. 1 | CD4$^+$ T cells (susbsets) | RNA-seq | Ranzani et al., 2015 | ERP004883 (ENA) |
| Fig. 1 | Cell lines | RNA-seq | Martinez et al., 2020 | GSE125218 (GEO) |
| Fig. 2 | Immune cells | MS-based proteomics | Rieckman et al., 2017 | Supplementary material |
| Fig. 2 | Cell lines | MS-based proteomics | Geiger et al., 2012 | Supplementary material |
| Fig. 3 | CD4$^+$ T cells | RNA-seq | Wolf et al., 2020 | GSE147229 (GEO) |
| Fig. 3 | CD4$^+$ T cells | MS-based proteomics | Wolf et al., 2020 | https://www.immunomics.ch |
| Fig. 3 | CD4$^+$ T cells | SILAC-derived protein synthesis rates | Wolf et al., 2020 | https://www.immunomics.ch |
| Fig. S8 | HeLa | Measured m7G sites | Zhang et al., 2019 | Supplementary material |
| Fig. S12 | 3'UTR library | DNA-seq | This manuscript | GSE240919 |

GEO: Gene Expression Omnibus of the National Center for Biotechnology Information (NCBI); ENA: European Nucleotide Archive;

**Data S1 to S5:**

**Data S1. SONAR feature importance from protein abundance models (separate file)**
Data related to Figure 1.

**Data S2. Cell-type specific SF usage of protein abundance models (separate file)**
Data related to Figure S3.

**Data S3. Syn3UTR sequences (separate file)**
Data related to Figure 3.

**Data S4. Massively parallel reporter assay motif enrichment in HEK293, $CD4^+$ and $CD8^+$ T cells (separate file)**
Data related to Figure 3.

**Data S5. SONAR feature importance from mRNA abundance, translational rate, protein abundance in $CD4^+$ T cells throughout activation (separate file)**
Data related to Figure 4, 5, S6.

**Data S6. Relation of mRNA abundance and putative m7G sites in HeLa cells (separate file)**
Data related to Figure S8.
